# A dirofilariasis mouse model for heartworm preclinical research

**DOI:** 10.1101/2023.04.03.535321

**Authors:** A E Marriott, J L Dagley, S Hegde, A Steven, C Fricks, U DiCosty, A Mansour, E J Campbell, C M Wilson, F Gusovsky, S A Ward, W D Hong, P O’Neill, A Moorhead, S McCall, J W McCall, M J Taylor, J D Turner

**Author notes:** Joint first authors (A E Marriott, J L Dagley).

## Abstract

Use of experimental cats and dogs in veterinary heartworm preclinical drug research is increasing. As a potential alternative primary *in vivo* heartworm preventative drug screen, we assessed lymphopenic mice with ablation of the interleukin-2/7 common gamma chain (γc) as susceptible to the larval development phase of *D. immitis*. Non-obese diabetic (NOD) Severe Combined ImmunoDeficient (SCID)γc^-/-^(NSG / NXG) mice consistently yielded viable *D. immitis* larvae at 2-4 weeks post-infection across multiple experiments, different batches of infectious larvae inoculates, different isolates of *D. immitis* and at independent laboratories. Mice did not display any overt clinical signs associated with infection up to 4 weeks. Developing larvae were found in subcutaneous and muscle fascia tissues, the natural site of this stage of heartworm in dogs. Larvae retrieved from NSG / NXG mice were mid-L4 stage of development. Compared with 14-day *in vitro* propagated larvae, *in vivo* derived L4 were significantly larger and contained expanded *Wolbachia* endobacteria titres, determined by QPCR and Fluorescent *in situ* Hybridisation (FISH). We established an *ex vivo* 6-day L4 paralytic screening system against nematodicidal agents (moxidectin, levamisole) which highlighted discrepancies in relative drug sensitivities in comparison with *in vitro* reared L4 *D. immitis.* We demonstrated effective depletion of *Wolbachia* by 70-90% in *D. immitis* L4 following 2-7 day oral *in vivo* exposures of NSG / NXG infected mice with doxycycline or the rapid-acting investigational anti-*Wolbachia* drug, AWZ1066S. We validated the NSG / NXG mouse model as a filaricide drug screen by *in vivo* treatments with single injections of moxidectin, which mediated 60-88% reduction in L4 larvae at 14-28 days. Future adoption of the mouse model as a first-line efficacy screen will benefit end-user laboratories conducting research and development of novel heartworm preventatives via increased access, rapid turnaround and reduced costs whilst simultaneously decreasing need for experimental cat or dog use.

## INTRODUCTION

*Dirofilaria immitis* is a major veterinary filarial parasitic nematode causing chronic heartworm disease (HWD) in dogs. Dirofilariasis is spread primarily by mosquito species of the *Culicidae* family, including the invasive tiger mosquito, *Aedes albopictus* [1]. HWD develops following the establishment of adult nematodes in the right chambers of the heart associated vessels following larval migrations in subcutaneous and muscle tissues. Adult infections can persist in the heart for greater than five years [2]. Pathology is chronic-progressive, associated with enlargement and hyper-proliferation of endocardium and physical blockage of adult worms in the pulmonary artery contributing to vessel narrowing, hypertension and ultimately heart failure [3]. *Dirofilaria immitis* causes a more acute immunopathology in cats where arrival of immature worms often triggers an overt inflammatory reaction in the lungs leading to heartworm-associated respiratory disease [2]. Both cats and dogs are at risk of acute, fatal thromboembolisms when dead adult worms lodge in pulmonary vasculature [3]. *Dirofilaria spp.* can also cause abbreviated zoonotic infections in humans, whereby arrested development of immature adults can cause subcutaneous nodules and lung parenchyma disease [4]. *Dirofilaria repens* is the most widely reported dirofilarial zoonosis, noted to be increasing across Europe, Asia and Sri Lanka; although, *D. immitis*, *D. striata, D. tenuis, D ursi* and *D. spectans* also infect humans [5]. In 2012, 48,000 dogs tested positive for heartworm in the USA and in 2016 over one million pets were estimated to carry disease [6]. Incidence of HWD in the US is increasing both within endemic areas and into erstwhile HW-free, westerly and northerly regions, including Canada [3]. A similar epidemiological pattern of increased dirofilariae incidence in The Mediterranean and spread into northern latitudes of Central and Western Europe has also been documented [7, 8].

Heartworm disease is controlled mainly by preventative chemotherapy and curative treatment of diagnosed cases. Chemo-prophylaxis with macrocyclic lactones (ML), namely: ivermectin, milbemycin oxime, moxidectin and selamectin, are effective at targeting L3-L4 larvae during subcutaneous tissue development and before immature adults reach the pulmonary artery to establish pathological adult infection [9, 10]. After more than 40 years of use in veterinary medicine, ML drug resistance is prevalent in veterinary nematode parasites, with several *D. immitis* isolates formally determined as resistant to ML, whereby timed experimental infections and accurate prophylactic dosing have failed to prevent the development of fecund adult HW infections [10].

The only regulatory approved cure available for HWD is the injectable, melarsomine dihydrochloride. However, issues of this therapy include: lengthy treatment regimens requiring in-clinic administrations, potential steroid pre-treatment, exercise restriction, and the risk of severe adverse events. Melarsomine is unsafe for use in cats, with no alternative curative therapies currently approved. Alternative curative therapies include the use of moxidectin and doxycycline (‘moxi-doxy’) [11] with the latter antibiotic validated as a curative drug targeting the filarial endosymbiont, *Wolbachia*, demonstratable in human filariasis clinical trials [12]. However, due to concerns of veterinary applications of doxycycline, in companion animals, such as long treatment timeframes, dysbiosis side-effects, and antibiotic stewardship of a human essential medicine, development of short-course, narrow-spectrum anti-*Wolbachia* heartworm therapeutics without general antibiotic properties, may offer a potential future alternative [13].

ML preventatives, costing typically between $266-329 a year for a pet’s treatment in the USA, represent a potential multi-billion dollar global market [14]. Due to emergent spread of *Dirofilaria immitis* infections, the growing concerns of ML prophylactic failure in USA and the current inadequacies of curative treatments, new therapeutic strategies are being intensively investigated. The only fully validated *in vivo* preclinical models currently available for heartworm anti-infectives are laboratory reared cats and dogs.

Lymphopenic and type-2 immunodeficient mice have been recently developed and validated as *in vivo* and *ex vivo* drug screens for the medically important filarial parasite genera; *Brugia, Onchocerca* and *Loa* [12, 15–17]. Here, we describe that lymphopenic immunodeficient mice with ablation of the interleukin-2/7 common gamma chain (gc) are susceptible to the initial tissue larval development phase of *D. immitis* and can be successfully utilised as a primary drug screening model for evaluation of direct-acting preventatives and anti-*Wolbachia* therapeutics.

## RESULTS

### Selection of a susceptible mouse model of tissue-phase heartworm infection

NSG and RAG2γc mouse strains were initially selected to investigate permissiveness to *D. immitis* tissue-phase larval infections, based on our previous success in establishing long-term infections of the related filarial species, *Brugia malayi, Loa loa* and *Onchocerca ochengi,* utilising lymphopenic immunodeficient mice [15, 17]. In these models, the additional knockout of the interleukin-2/7 common gamma chain within lymphopenic mice is essential for susceptibility to *Loa* adult development in subcutaneous tissues [17] and bolsters both *Brugia* and *Onchocerca* adult infections within the peritoneal cavity [18]. We also trialled methylprednisolone acetate (MPA) administrations to evaluate whether steroid suppression of residual innate immune responses could increase survival and yields of *D. immitis* larvae *in vivo,* as has been reported for experimental *Strongyloides stercoralis* infections [19]. Initially, we used a Missouri (MO) isolate of *D. immitis* (NR-48907, provided by the NIH/NIAID Filariasis Research Reagent Resource Center, FR3, for distribution through BEI Resources). Infectious L3 were isolated 15 days after membrane feeding of *D. immitis* mf in dog blood to *A. aegypti* (Figure 1A). Following inoculations of 200 L3 under the skin, at 14 days post-infection, we successfully recovered *D. immitis* parasites from subcutaneous and muscle fascia tissues in all (5/5) NSG and RAG2γc+MPA mice (Figure 1B). Multiple tissues were dissected to locate parasites (heart, lungs, peritoneal cavity, gastrointestinal tract, liver, spleen) but no evidence of infection was found in these ectopic locations. Infection success was lower in NSG+MPA (3/4 mice) and RAG2γc (4/5) mouse groups. Yields significantly varied between groups with RAG2γc+MPA mice yielding higher numbers of *D. immitis* developing larvae (L4) compared with either RAG2γc or NSG+MPA groups (Kruskal-Wallis 1 Way ANOVA P=0.0033, Dunn’s post-hoc tests P<0.05). Median recovery rates were similar between NSG and RAG2γc+MPA groups (median recovery 9% vs 12%, ns).

**Figure 1.**
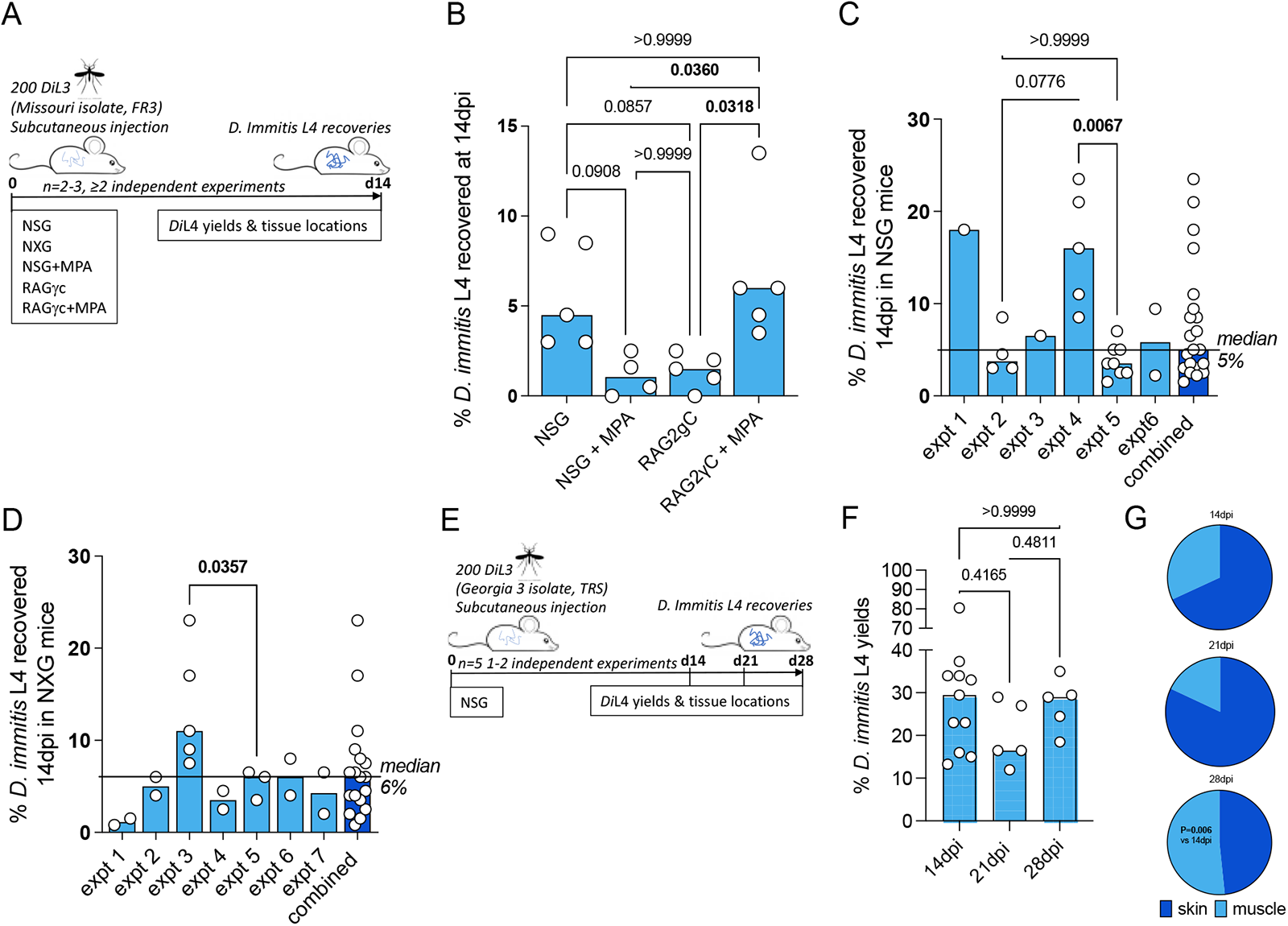
Susceptibility of compound immunodeficient mouse strains to D. immitis. Experimental schematic using Missouri (MO) isolate D. immitis (A). Percentage recovery of initial infectious load of D. immitis L4 at 14 days post-infection in indicated immunodeficient mice (B-D). Experimental schematic using Georgia (GA)-3 isolate D. immitis (E). Percentage recovery of initial infectious load (F) and tissue distributions (G) of D. immitis L4 at indicated time-points in NSG mice. Bars indicate median values with individual mouse data plotted. Significant differences were assessed by Kruskal Wallis One-way ANOVA with Dunn’s multiple comparison’s tests except (G) where difference in proportions were tested by Fisher’s Exact Test. Significant differences (P values <0.05) are indicated in bold. Data is combined of two or more independent experiments (B-D) or individual experiments (C,D,F) with between 1-5 mice per group.

Due to the simpler infection regimen in NSG mice, the international commercial availability of the model, the potential for further humanisation, and to avoid potential welfare or drug-drug interactions arising with long-term MPA administrations, we selected this immunodeficient mouse model for extensive characterisation. We evaluated the infection success and yields of *D. immitis* L4 across multiple independent experiments utilising different batches of MO isolate *D. immitis* shipped from USA to UK as mf in dog blood and passaged to the infectious L3 stage in *A. aegypti* mosquitoes (Figure 1C). In six independent experiments, using a total of 21 mice, we were reproducibly able to recover *D. immitis* larvae at 14dpi (21/21 mice) with a 5% median yield of the initial 200 L3 infectious inoculate (range 1.5% – 23.5%). In experiments where >2 mice were infected, we compared yields between batches and determined that batch-to-batch variability of the initial L3 infectious inoculate significantly influenced the yields of larvae recovered at 14dpi in NSG mice (Kruskal-Wallis 1Way ANOVA P=0.0017, Dunn’s post-hoc tests P<0.01). We then measured recoveries utilising NXG mice; a similar Severe Combined Immunodeficient mouse line on the Non-Obese Diabetic strain background with additional γc ablation, developed independently and recently commercialised by Janvier Laboratories [20] (Figure 1D). As with the original NSG mouse line, in seven independent experiments, with a total of 18 experimental mouse infections, we consistently recovered developing larvae at 14dpi (18/18 NXG mice) with a 6% median proportion of the initial 200 L3 infectious inoculate (range 1% – 23%). Similar to NSG infections, in individual experiments where >2 NXG mice were available for analysis, batch-to-batch variability of the initial L3 infectious inoculate significantly influenced the yields of larvae recovered at 14dpi (Mann-Whitney P<0.05). We then repeated experiments with NSG mice in TRS Labs (Georgia, USA), accessing an in-house parasite life-cycle and using a unique ‘Georgia III’ (GA III) isolate of *D. immitis*. We infected batches of five mice and evaluated yields of L4 at 14, 21 or 28 days post inoculation with 200 L3 (Figure 1E). All mice, irrespective of time point post-inoculation, yielded GA III *D. immitis* developing larvae. Yields were typically 6-fold higher than those derived at the LSTM laboratory at 14 days post-infection (median = 29.5%, range 13.3-80.5%, Figure 1F). Yields did not significantly deviate between 14, 21 and 28 days post-infection (Figure 1F). However, the distribution of larvae in mouse tissues had changed between 14 and 28 days-post infection, with relatively more larvae recovered in muscle tissues by 28 dpi (P<0.01, Fisher’s Exact Test, Figure 1G). These experiments demonstrate the reproducible success of NSG/NXG mouse models as susceptible to *D. immitis* tissue-stage infection using different isolates of heartworm, in independent laboratories and when shipping larvae internationally between sites. Further, time-course data indicates that tissue-phase heartworm larvae persist without significant decline in yields within NSG mice whilst initiating their natural migratory route through subcutaneous and muscle tissues over the first 28 days of infection.

NSG mouse-derived developing larvae demonstrate superior morphogenesis, *Wolbachia* content and reduced drug assay sensitivities compared with *in vitro* cultured *D. immitis*

Mosquito-derived infectious L3 larvae are traditionally utilised in serum-supplemented 37°C mammalian cultures to induce moulting and morphogenesis into fourth-stage developing larvae [21–23]. This technique has been utilised to study *D. immitis* larval biology and for applied applications such as biomarker and preventative drug discovery [24–26]. The survival of various filarial parasite life-cycle stages can be extended when utilising co-cultures with mammalian ‘feeder cell’ monolayers or trans-well compartments [16, 27–31]. We therefore compared survival and motility of MO isolate *D. immitis* larvae between cell-free and LL-MCK2 (monkey) or MDCK (dog) kidney cell co-cultures in 10% calf-serum cultures (Figure 2A). The 50% survival time of cell-free cultures was d18 and subsequently, all larvae had died by d28 in culture (Figure 2B). Conversely, co-cultures with both LL-MCK2 and MDCK cells significantly increased survival, whereby >80% of *D. immitis* larvae were viable up to d28 (P<0.0001, Mantel Cox survival analysis). We noted a reduction in motility in all larval cultures after the first week in culture, which persisted to end-point, apart from MDCK co-cultures which returned to full-motility by d32 in culture (Supplementary figure 1). Selecting MDCK co-cultures as supportive of long-term larval motility and survival, we directly compared morphogenesis, growth and *Wolbachia* endobacterial expansions between *in vitro* propagated MO *D. immitis* larvae and MO larvae derived from NSG mouse infections at the 14-day time point (Figure 2A). Both *in vitro* and *in vivo* derived d14 *D. immitis* larvae displayed the blunted and widened anterior extremities characteristic of the L4 developmental stage [32, 33], compared with the tapered, narrow anterior of filariform infectious L3 (Figure 2C). However, anterior morphogenesis was partially arrested in *in vitro* compared with NSG mouse-derived larvae (Figure 2C). In the dog, larvae complete the L3-L4 moult rapidly, the vast majority by 3 days post-infection [34]. In our cultures, approximately 50% of the day 14 L4 had completed moulting, with cuticle casts evident in the culture media. The other ∼50% of *in vitro* cultured larvae displayed partial moulting of the third stage cuticle (Figure 2C). There were obvious microscopic degenerative features of the *in vitro* larvae by d14 compared with *in vivo* larvae, including malformed cuticle, hypodermis, buccal cavity, oesophagus and intestine (Figure 2C). Despite their high survival rate and continued motility, 14-day old *in vitro* propagated larvae were also significantly stunted compared with larvae derived from NSG mice (mean = 1020 *vs* 1880 μM, 1 way ANOVA F=57.7, P<0.0001, Tukey’s multiple comparisons test), and had not grown significantly in comparison to the L3 infectious stage (mean = 870 μm) (Figure 2D). *Wolbachia* titre analysis by QPCR further highlighted disparities between *in vitro* and *in vivo* reared larvae (Figure 2E). The MO *D. immitis in vivo* larvae had undergone a significant, 66-fold average *Wolbachia* expansion during the 14-day NSG mouse infection in comparison to iL3 (median = 4.2×10^4^ *vs* 6.2×10^2^ *Wolbachia*/larva, Kruskal Wallis 1-way ANOVA 26.4, P<0.0001 Dunn’s multiple comparisons test), whereas MO larvae cultured for 14 days *in vitro* had failed to expand *Wolbachia* content (median = 8.7×10^2^ *Wolbachia*/larva).

**Figure 2.**
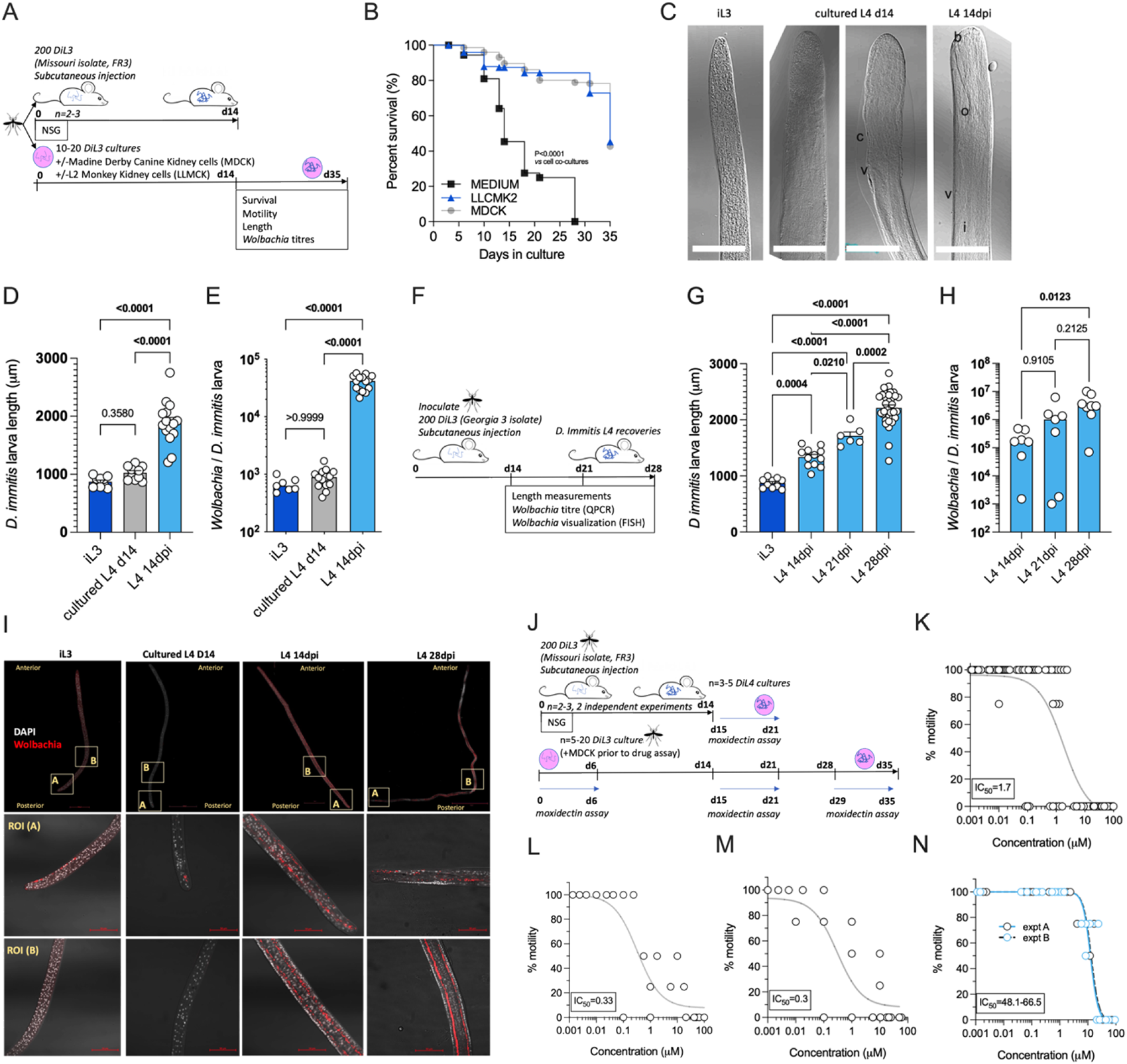
comparative morphogenesis, Wolbachia expansions and drug assay sensitivities of NSG-mouse derived larvae compared with in vitro cultured D. immitis. Experimental schematic using MO isolate D. immitis L3 (A). Survival analysis of cultured larvae in indicated conditions (B) representative photomicrographs of iL3, d14 L4 cultures or L4 recovered from NSG mice at 14dpi (C; b=buccal cavity, c=cuticle, i=intestine, o=oesophagus, v=vulva, scale bars = 50μM). Length (D) and Wolbachia content (E) of iL3, d14 L4 cultures or L4 recovered from NSG mice at 14dpi. Experimental schematic using GA3 isolate D. immitis L3 (F). Length (G) and Wolbachia content (H) of NSG mouse derived L4 larvae at indicated time-points. Representative FISH photomicrographs of iL3, d14 L4 cultures or L4 recovered from NSG mice at 14 or 28dpi (Wolbachia 16SrRNA red, DAPI grey, scale bars = 50μM). Experimental schematic of drug assays utilising MO isolate D. immitis cultures or ex vivo larvae from NSG mice (J). Moxidectin concentration D. immitis motility inhibition analysis after six days exposure when using cultured D. immitis for periods between 0-6d L3/L4 (K) 15-21d L4 (L) 28-35d L4 (M) or 15-21d L4 derived from 14dpi NSG mice (N). In K-N, non-linear curves are 3-parameter least squares fit with [IC_50_] calculated in Prism 9.1.2. Bars represent mean +/-SEM (D, G) or median (E, H) values with individual larva data plotted. Significant differences were determined by Mantel Cox log-rank tests (B) One-way ANOVA with Tukey’s multiple comparisons tests (D, G), or Kruskal Wallis with Dunn’s multiple comparisons tests (E, H). Significant differences (P values <0.05) are indicated in bold. Data is one individual experiment except (N) which is two independent experiments.

Utilising GA III *D. immitis,* we further examined length and *Wolbachia* expansions between d14 and d28 post-infection in NSG mice (Figure 2F). GA3 L4 continued to grow in length between d14, d21 and d28 post-infection in NSG mice (Figure 2G; means = 1335, 1713 & 2211μm, respectively, 1 Way ANOVA F=87.4, P<0.05 – P<0.0001, Tukey’s multiple comparisons tests). Similarly, *Wolbachia* titres continued to expand within the NSG-derived GA III *D. immitis* larvae (Figure 2H) with significant differences evident between d14 and d28 (median = 1.7×10^5^ *vs* 3.1×10^6^ *Wolbachia/*larva, Kruskal Wallis statistic = 8.5, P<0.05 Dunn’s multiple comparisons test). We corroborated quantitative PCR *Wolbachia* data, visualising that time-dependent *Wolbachia* multiplication was occurring within the hypodermal chord cell syncytia from a posterior to anterior direction in NSG mouse-derived, but not *in vitro* cultured, *D. immitis* L4 specimens, utilising fluorescent *in situ* hybridisation (FISH) of *Wolbachia* 16S rRNA and confocal microscopy (Figure 2I). We then examined the *in vitro* vs *ex vivo* paralytic susceptibilities of MO isolate *D. immitis* L4 following 6-day exposures to the standard preventative drug, moxidectin, using cultured L3-L4 larvae at 0-6d, 15-21d or 28-35d compared with NSG mouse L4 larvae isolated at 14dpi and exposed to drug *ex vivo* between 15-21d in matching culture conditions (Figure 2J). The IC_50_ concentrations inhibiting motility of *D. immitis* were 1.7μM for 0-6d L3-L4 larvae (Figure 2K). Sensitivity to moxidectin had increased in d14-d35 larvae with IC_50_ ranging between 300-330nM (Figure 2L&M). In comparison, *ex vivo* larvae derived from NSG mice were relatively insensitive to the *in vitro* paralytic activity of moxidectin with IC_50_ ranging between 48-66μM (Figure 2N). This equated to a >28-fold decrease in moxidectin susceptibility compared with *D. immitis* L3-L4 cultures and >140-fold decreased sensitivity compared with longer-term L4 cultures. We further examined relative paralytic susceptibilities of *D. immitis* MO *in vitro* vs *ex vivo* L4 to 6-day exposures of the anthelmintic, levamisole, commencing at 15 days after iL3 culture / infection (Supplementary Figure 2). Whilst *in vitro* larvae were susceptible to high doses of levamisole (IC_50_ 13.2 μM), *ex vivo* L4 maintained full motility for 6 days in the presence of the top dose of drug (100 μM). These data demonstrate the developmental superiority of *D. immitis* larvae derived from the subcutaneous and muscle tissues of NSG mice compared with standard *in vitro* cultures, reflected in a profound lowered sensitivity to direct-acting nematodicidal agents when used in *ex vivo* drug titration assays.

### *D. immitis* NSG mouse infections can be used to evaluate anti-*Wolbachia* drug efficacy

Because we established rapid *Wolbachia* expansions occur during the L4 tissue development phase of *D. immitis* following infections of NSG mice, we next investigated the validity of the *D. immitis* NSG mouse model as an anti-*Wolbachia* drug screening model. We infected batches of 4-6 mice with MO isolate *D. immitis* iL3 and randomised mice into 7-day 50 mg/kg oral treatment with doxycycline or matching vehicle controls, commencing at infection with a further 7-day washout period to 14dpi (Figure 3A). We selected this regimen and timing of dose based on proven significant depletion of *B. malayi* L3-L4 *Wolbachia in vivo* in a SCID mouse model [35]. We then randomised a further four mice into a 2-day bi-daily 200mg/kg treatment of our fast-acting anti-*Wolbachia* azaquinazoline clinical candidate, AWZ1066S [36] or vehicle control, to compare relative anti-*Wolbachia* activity. Doxycycline treatment mediated a 70% median reduction in *Wolbachia* titres in d14 MO *D. immitis* larvae when compared against vehicle control levels (0.29×10^4^ vs 9.5×10^4^ *Wolbachia*/larva, Mann-Whitney test P=0.014, Figure 3B). The short-course AWZ1066S 2-day oral treatment mediated a more profound 90% median efficacy in depletion of *Wolbachia* from d14 MO *D. immitis* larvae (0.49×10^4^ vs 4.5×10^4-^ *Wolbachia*/larva, Mann-Whitney test P<0.0001, Figure 3C). The effect of *Wolbachia* depletion via doxycycline and AWZ1066S on larval growth was also evaluated (Figure 3D&E). We found that depletions of *Wolbachia* by 7-day doxycycline or 2-day AWZ1066S were associated with a 15.9% and 15.3% mean stunting effect on 14d MO *D. immitis* larvae, which was significant for doxycycline treatment (Student’s T-Test, P=0.0182). We repeated the validation of the *D. immitis* NSG mouse model as an anti-*Wolbachia* drug screening system in an independent laboratory, utilising GA III isolate *D. immitis* (Figure 3A). In this dosing study, the 7-day oral regimen of doxycycline mediated a significant, 89% median depletion of *Wolbachia* in d14 GA3 larvae (0.25×10^5^ vs 2.4×10^5^ *Wolbachia*/larva, Mann-Whitney test, P=0.0098, Figure 3F). We corroborated the clearance of *Wolbachia* from posterior hypodermal chord cells by FISH staining (Figure 3G). Stunting of GA III *D. immitis* larvae was also apparent following 7-day doxycycline exposures in NSG mice (mean reduction in length 30%, Figure 3G). Together, these data demonstrate the utility of the *D. immitis* tissue-phase NSG mouse model to screen for efficacy of oral anti-*Wolbachia* regimens *in vivo*.

**Figure 3:**
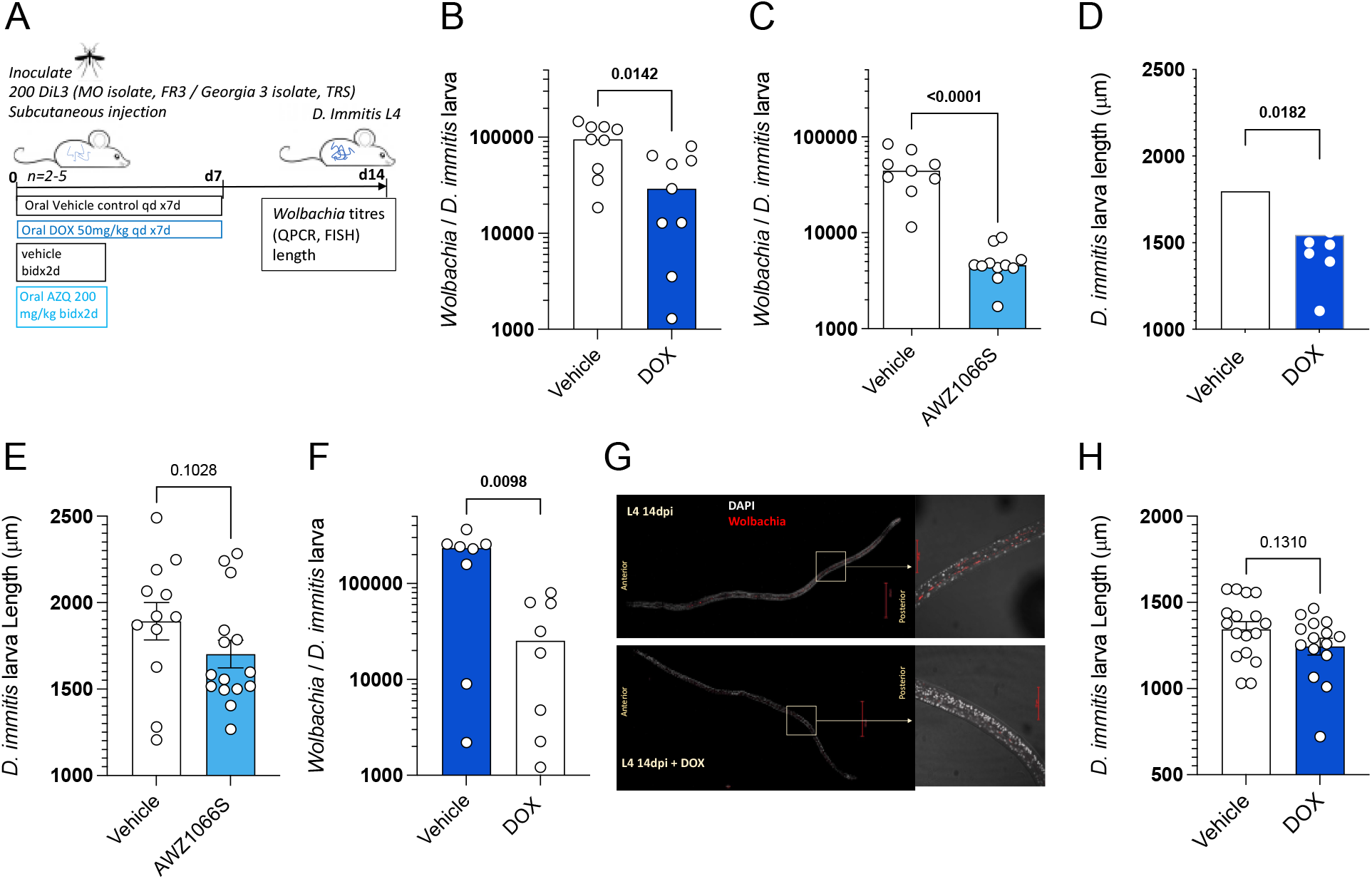
validation of the D. immitis NSG mouse model as an in vivo anti-Wolbachia drug screening system. Experimental schematic using MO or GA III isolates of D. immitis L3 (A). Wolbachia loads determined by PQCR in d14 MO D. immitis larvae exposed to doxycycline (B) or AWZ1066S (C). Length changes in d14 MO D. immitis larvae exposed to doxycycline (D) or AWZ1066S (E). Wolbachia loads determined by QPCR in d14 GA III D. immitis larvae exposed to doxycycline (F). Visualisation of Wolbachia depletion in hypodermis of d14 GA3 D. immitis larvae exposed to doxycycline by FISH (G). Length changes in d14 GA III D. immitis larvae exposed to doxycycline (H). Bars represent median (B, C, F) or mean +/-SEM (D, E, H) values with individual larva data plotted. Significant differences were determined by Mann-Whitney tests (B, C, F) or unpaired Student’s T Tests (D, E, H). Significant differences (P values <0.05) are indicated in bold. Data is one individual experiment with larvae derived from 2-5 mice per group.

### *D. immitis* NSG mouse infections can be used to evaluate preventative drug efficacy

Moxidectin is a front-line ML preventative used in various oral, topical or injectable formulations as monthly, biannual or annual heartworm prophylaxis in dogs [37]. We selected a single high dose subcutaneous injection of moxidectin (2.5mg/kg) for evaluation of larvicidal efficacy in NSG/NXG mice (emulating route of delivery and dose of long-acting injectable formulations of moxidectin in dogs). Matched pairs of mice were infected with batches of 200 *D. immitis* MO or GA III isolate larvae and the next day were randomised into vehicle control or moxidectin treatment (Figure 4A). After 14 days post infection (13 days post treatment), we recorded 65-80% reductions in MO *D. immitis* L4 in NSG mice (n=3 pairs, P=0.03, paired t-test, Figure 4B). A similar range of larvicidal efficacy were evident in NXG mice 14 days after infection with MO isolate *D. immitis* and treatment with a single injection of moxidectin (range 46-88% n=6 pairs, P=0.008, Figure 4C). When using the GA III isolate of *D. immits* for NSG infections, level of moxidectin efficacy ranged between 29-73%, evaluated at 14 days post-infection (P=0.022, n=5 pairs, Figure 4D). We examined extended washout periods after moxidectin single dosing in NSG mice infected with GA III *D. immitis*. At 21 days post infection, the range of moxidectin efficacy was 45-94% (P=0.032, n=5 pairs, Figure 4E) whilst at 28 days post-infection, efficacy ranged between 35-91% (P=0.0078, n=5 pairs, Figure 4F). The median efficacy for all studies is shown in Figure 4G. In summary, single injection of moxidectin delivered a median efficacy between 65-80% at two weeks in NSG/NXG mice infected with MO isolate, and 60, 73 and 75% efficacy at two, three or four weeks in NSG infected with GA III isolate. All mice on drug studies displayed typical behaviour and gained weight over the 2-4 week period of infection and dosing (Supplementary figure 3).

**Figure 4:**
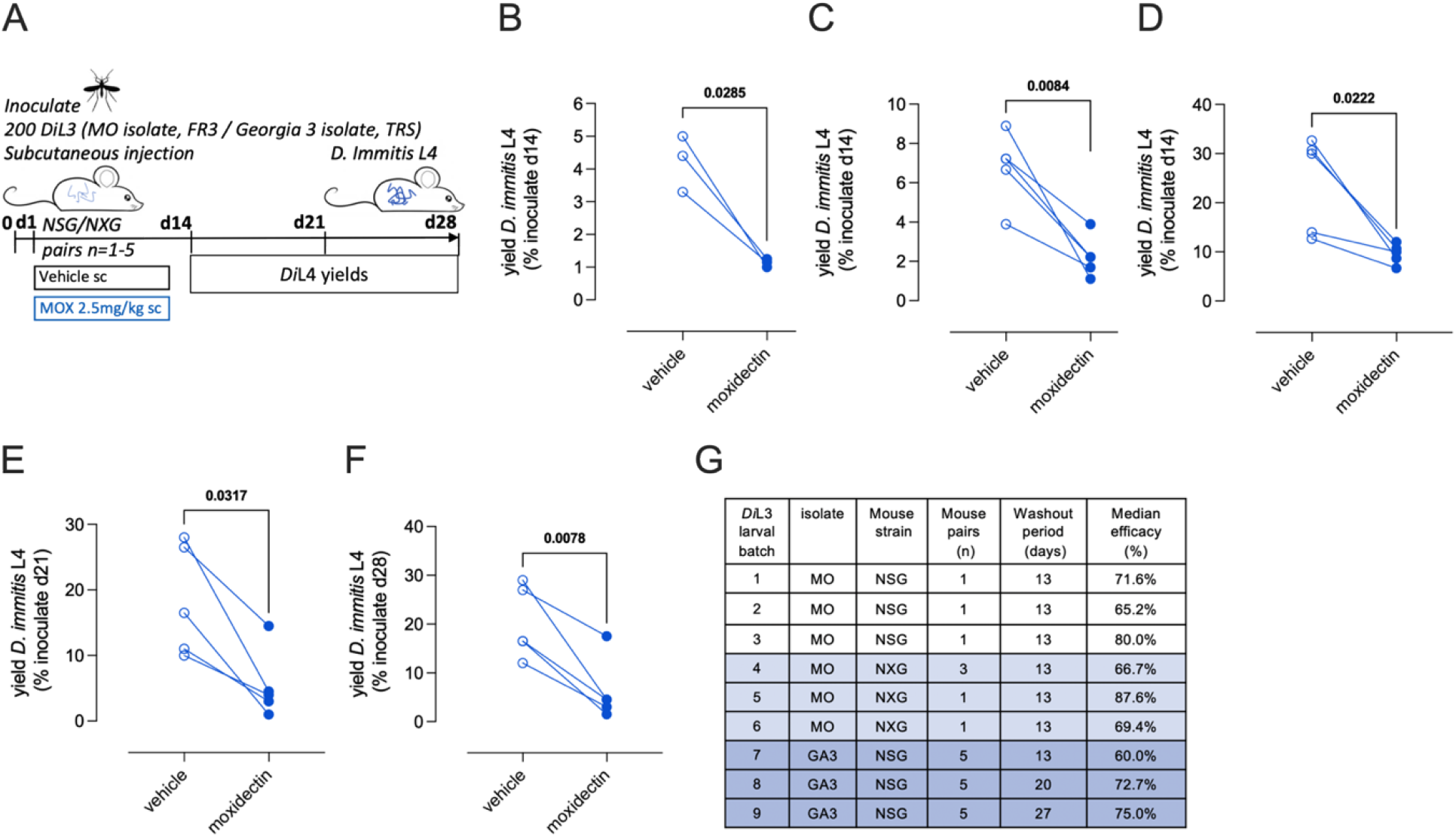
Validation of D. immitis NSG mouse model as an in vivo larvicidal drug screen. Experimental schematic using MO or GA III isolates of D. immitis L3 (A). Change in D. immitis larval recoveries at d14 in randomised NSG (B) or NXG (C) mouse pairs inoculated with MO isolate D. immitis and treated with vehicle or 2.5mg/kg moxidectin via subcutaneous injection at d1. Change in D. immitis larval recoveries at d14 (D) d21 (E) or d28 (F) in randomised NSG mouse pairs inoculated with GA III isolate D. immitis and treated with vehicle or 2.5mg/kg moxidectin via subcutaneous injection at d1. Summary of median moxidectin efficacies derived from NSG/NXG mouse pairs inoculated with different batches of MO/GA3 L3 D. immitis larvae and treated with vehicle or moxidectin, evaluated at 14, 21 or 28 days. Significant differences were determined by paired T-tests. Significant differences (P values <0.05) are indicated in bold. Data is pooled from three individual experiments (B, C) or one individual experiment (D-F), derived from 1, 3 or 5 mouse pairs.

## DISCUSSION

We have determined that ablation of both the B & T lymphocyte compartment and additional cytokine signalling via the IL-2/7 common gamma chain receptor in mice allows permissiveness to *D. immitis* tissue phase larval development over the first 28 days of infection. A polarised type-2 adaptive immune response with associated type-2 tissue macrophage activation leads to eosinophil entrapment and degranulation as the basis of immune-mediated filarial larvicidal activity in mice [38–41]. However, experimental infections with *B. malayi, L. sigmodontis* and *O. ochengi* in lymphopenic mouse strains (SCID / RAG2^-/-^) with additive γc gene ablations have illustrated bolstered chronic susceptibility [18, 42], whilst in *L. loa* subcutaneous infections, only combination of lymphopenia and γc deficiency is sufficient to allow permissiveness to adult infections [43]. Thus, an additional layer of innate immune resistance operates which can eliminate establishing larval filarial infections. In *B. malayi* infections, p46+ NK cells with an activated / memory phenotype and residual eosinophilia are implicated in the innate immune resistance to chronic infection in RAG2^-/-^mice [18] whereas in *Litomosoides* infections, CD45-/TCRβ-/CD90.2+/Sca-1+/IL-33R+/GATA-3+ type-2 innate lymphoid cells (ILC2) are required for innate immune resistance to microfilarial blood infections [44]. It remains to be determined which of these multiple innate and adaptive immune processes are operating to control larval establishment of *D. immitis* in mice. Because we detected increases in *D. immitis* larval burdens in NOD.SCID-vs BALB/c RAG2^-/-^-γc^-/-^ mice which could be improved by steroid treatments in the latter model, this may indicate residual innate immune differences between background strains. For instance, NOD mice are deficient in complement humoral immunity due to a 2-bp deletion in the hemolytic complement (Hc) gene, which encodes the C5 complement protein [45]. Our model now affords an opportunity for reconstitution of innate or adaptive immune cell types and humoral immune components to dissect the mechanisms of immunity to *D. immitis* migrating larvae, as has recently been attempted for *L. sigmodontis* with CD4+ T cell transfers into RAG2^-/-^γc^-/-^ mice [46]. This application of the model may be useful in determining minimally sufficient immune pathways necessary to mediate sterilising immunity. Similarly, the model may be valuable in evaluating efficacy of neutralising sub-unit vaccine target antibody responses (e.g., via passive transfer of purified specific antibodies or isolated B cell clones adoptively transferred from immunocompetent NOD mice).

We evaluated that *D. immitis* larval parasitism and development in our mouse model accurately tracts the natural course of infection in definitive hosts over the first month. All larvae were recovered from the subcutaneous tissues and muscle fascia, in line with previous observations of natural parasite locations in both ferret and dog infections of *D. immitis* at this time-interval [33, 47]. We demonstrate that *in vivo* larvae complete cuticle moulting and undergo 4^th^ stage larval morphogenesis. L4 growth lengths in NSG/NXG mice are also within the range of those prior documented in dog and ferret infections at matching point of infection, at 14-15 days (NSG = 1.2-2.8 mm, dog = 1.7-2.2 mm, ferret = 1.6–2.7 mm) [33, 47].

We also demonstrate that *D. immitis* expand *Wolbachia* titres significantly during parasitism of NSG mice. From PCR analysis, we ascertain that *Wolbachia* are doubling approximately every 42h for MO isolate to every 55h for GA III isolate over the first 14-day infection time-course of *D. immitis* L3-L4 larvae *in vivo*. This is the first record of early *Wolbachia* expansions in *D. immitis* developing larvae and is comparatively slower compared with the average doubling time (32h) over the first 14 days of L3-L4 development *in vivo* for the human filariae, *B. malayi* [48]. The establishment of the *D. immitis* mouse model now allows for tractable comparative endosymbiotic biology of this clade C nematode *Wolbachia* (also found in the causative agent of river blindness, *O. volvulus*) versus the clade D *Wolbachia* of human lymphatic filariae, most commonly used in basic and applied nematode *Wolbachia* research.

Until now, a ready source of *in vivo D. immitis* L4 propagations for onward *‘ex vivo*’ basic and translational research has been unavailable. Mosquito stage L3 can be induced to moult rapidly into the early L4 stage, with as much as 95% moulting success, and survive for 3 weeks in calf-serum supplemented cultures [21]. We recapitulated this early L4 morphogenesis and improved L4 longevity to greater than one month in culture if larvae were co-cultured with dog or monkey kidney cells. However, comparisons with *in vivo* reared larvae highlighted several defects in growth, incomplete morphogenesis and, most strikingly, an almost complete failure to expand *Wolbachia* endosymbiont titres. Therefore, the failure of larvae to thrive *in vitro* may be linked with a deficit in *Wolbachia-*produced haem, riboflavin, nucleotides, or other biosynthetic pathways identified as relevant in the *Wolbachia –* nematode symbiosis [49]. Whether environmental cues are lacking *in vitro* for *Wolbachia* expansion is currently not known. Sub-optimal neo-glucogenesis in cultured *D. immitis* larvae could lower available carbon energy sources necessary for *Wolbachia* expansion [50]. Alternatively, because autophagic induction in filariae regulates *Wolbachia* populations residing within host vacuoles [51], failure of *Wolbachia* growth may be the result of starvation / stress in culture inducing autophagy. Certainly, filarial stress responses are demonstrably upregulated in *ex vivo* adult worm culture systems [52].

We exemplify that use of sub-optimal L3/L4 grown *in vitro* for pharmacological screening leads to significantly >28-fold increased sensitivities to the paralytic activities of two nematodicical agents, moxidectin and levamisole, compared with larvae of same age derived from NSG/NXG mouse infections. Thus, we argue that reliance on *in vitro* larvae may lead to spuriously artificial sensitivities to new preventatives in development and, onwards, to incorrect selection of candidates or dose levels for *in vivo* preclinical evaluations with consequences for incorrect cat and dog usage. The new mouse model now affords a facile method of generating physiologically relevant L4 larvae for more accurate pharmacological assessments prior to a decision to advance into *in vivo* preclinical screening justifying protected animal use.

Our data determining the failure of *Wolbachia* to expand *in vitro* within *D. immitis* developing larvae precludes use of cultured L3/L4 in evaluating activities of novel anti-*Wolbachia* compounds. We thus demonstrate utility of the NSG/NXG mouse model as an *in vivo* anti-*Wolbachia* drug screen. L4 larvae could be reproducibly depleted of *Wolbachia* using a seven-day regimen of the established anti-*Wolbachia* antibiotic, doxycycline, with confirmatory experiments run in an independent laboratory with a different *D. immitis* isolate. Excitingly, we demonstrate that a two-day *in vivo* treatment with the novel investigational azaquinazoline drug, AWZ1066S, is a rapid and profound *D. immitis Wolbachia* depleting agent with 90% efficacy achieved. This benchmark of 90% efficacy has been determined as clinically relevant in terms of sustained *Wolbachia* reductions and subsequent long-term anti-parasitic activities in human filariasis clinical trials [12]. The unique rapid activity of AWZ1066S has been determined through time-kill assays with *B. malayi Wolbachia,* whereby a near maximum kill rate can be achieved with 1-day exposure compared with six days for standard classes of antibiotics including tetracyclines [36]. Thus, azaquinazolines or other novel anti-*Wolbachia* chemistry with similar rapid killing activity, as identified in high throughput industrial screening [53], might hold promise as new heartworm preventative or curative candidates and now can be triaged for activity utilising our novel *D. immitis* NSG/NXG mouse model.

We used a high single parenteral dose of moxidectin, mimicking extended-release formulations used in dogs [37], to evaluate the *D. immitis* NSG/NXG mouse model as a preventative drug screen. We demonstrated, using multiple batches of different ML-susceptible *D. immitis* isolates, in different evaluating laboratories, that injected moxidectin mediated significant 65-89% reductions in larvae assessed between 13-27 days post-exposure. Whilst drafting this manuscript, an independent study was published reporting a similar success of establishing *D. immitis* subcutaneous larval infections in NSG mice [54]. Hess and colleagues also measured ivermectin and moxidectin responses in infected NSG mice following oral doses ranging between 0.001-3mg/kg given on day 0, 15 and 30 post-infection. Their studies determine that the MO isolate and an ivermectin resistant JYD-34 isolate of *D. immitis* were equally sensitive to moxidectin with high but incomplete efficacy demonstrable after 0.01mg/kg dosing. They also show high levels of ivermectin efficacy against the MO, but not JYD-34 isolate, in dose titrations ranging between 0.01-3mg/kg. Thus, we conclude the *D. immitis* NSG/NXG model is robustly validated by multiple independent laboratories as a screening tool for assessing direct-acting nematodicidal agents over at least a 28-day infection window. Our *ex vivo* and *in vivo* drug response evaluations of *D. immitis* L3/L4 in NSG/NXG mice demonstrate the flexibility to establish this model in independent laboratories with different commercially available NSG/NXG lines. Further, we determine feasibility of international shipping of live mf in dog blood to produce L3s for onward experimental infections in NSG/NXG mice. Mouse infections utilising shipped L3s may allow for increased accessibility to this model in order to expand experimental *D. immitis* research out of the few specialist reference centres which maintain the full life cycle of the parasite and/or the mosquito vector.

Main limitations of our study are: 1) lack of data on full permissiveness to adult infections within murine cardiopulmonary vasculature, 2) lack of validation within female NSG/NXG mice and 3) less than 100% achievable moxidectin preventative efficacy response. In Hess *et al*, evaluation periods were extended and, whilst larvae continued to grow and mature within NSG mice, a divergence in growth compared with comparative dog studies was apparent after the first month of infection. Further, there was no evidence of immature adults arriving in the heart and lungs by 15 weeks [54]. The authors conclude either physiological or anatomical deficiencies may prevent the full development of *D. immitis* in NSG mice. However, full development of the highly-related subcutaneous filaria, *Loa loa,* is possible in NSG mice [43] and thus it remains to be tested whether full *D. immitis* development may be achieved over an extended time-frame. We selected use of male mice due to observations that even in immunodeficient systems [55], as well as in outbred gerbils [56], male biased sex-specific susceptibility is a feature of rodent filarial infections. For future pharmacological investigations, to fully enable characterisation of interactions between sex and the drug pharmacokinetic-pharmacodynamic (PK-PD) relationship, it would be useful to assess whether *D. immitis* are able to develop within female NSG mice. In natural hosts, injectable formulations of moxidectin are proven to mediate 100% preventative efficacies [37].The substantial yet incomplete moxidectin responses in our NSG/NXG model may reflect full efficacy evolves over an extended time-period, particularly considering that this ML depots in fatty subcutaneous tissues and delivers a long-tail of systemic exposure, detectable over one month [57, 58]. Alternatively, an immunopharmacological mode-of-action involving disruption of immunosuppressive parasite secretions and an activated host-immune response has been proffered as one rationale why filarial larvae are differentially sensitive to ML-drugs at physiological levels *in vitro* vs *in vivo* [59]. Thus, we currently cannot rule out a potential synergy with adaptive immune-mediated responses (such as the development of opsonising or neutralising antibodies) contributing to the complete efficacy of moxidectin. As previously discussed, passive transfer of antibodies into NSG/NXG mice may determine whether such a mechanism contributes to ML preventative efficacy at physiologically relevant dose levels.

It is widely accepted that the use of specially protected, highly sentient species in preclinical research, including cats and dogs, should be strictly minimised wherever possible. This has not been plausible for veterinary heartworm preventative R&D due to a lack of a tractable small animal laboratory model. Typical drug screening has relied on *in vitro* potency testing against *D. immitis* larvae, potentially combined with initial preclinical evaluation in a surrogate rodent filarial infection model, before deciding to proceed into experimental dog infection challenge studies. Vulnerabilities of this approach include: differential drug sensitivities between larvae being tested *in vitro* versus *in vivo,* vagaries in filarial species larval migration routes/ parasitic niches, and variability in drug target expression / essentiality across different filarial parasite species and life-cycle stages, all of which may drive artefactual efficacy information. Our model, with international commercial supply of NSG/NXG strains and shipping of mf or L3 from donating laboratories, provides universal access to accurate and facile PK-PD assessments of preventative *D. immitis* drug candidate responses against the prophylactic L3-L4 larval target. Evaluations of drug larvicidal activities over the first month of infection, whilst larvae are developing in subcutaneous and muscle tissue, allows for rapid assessments whilst avoiding risk of welfare issues associated with arrival of adult parasites in the cardiovascular system. We observed no overt welfare issues of mice after parasitism, with mice gaining weight and displaying typical behaviour. If adopted, our model would accelerate drug research timelines and enable precise dose-fractionation studies for clinical selection.

We conclude that a *D. immitis* NSG/NXG mouse model is established for more efficient heartworm drug discovery which will reduce the requirements for long-term cat and dog experimentation with the risk to cause severe harm, in line with an ethos of ‘replacement, refinement and reduction’ of animals in scientific research.

## MATERIALS AND METHODS

### Animals

Male NOD.SCIDγc^−/−^ (NSG; NOD.Cg-*Prkdc^scid^ Il2rg^tm1Wjl^*/SzJ) and BALB/c RAG2^−/−^γc^−/−^ (RAG2ɣc; C;129S4-*Rag2^tm1.1Flv^ Il2rg^tm1.1Flv^*/J) mice were purchased from Charles River UK. Male NXG mice (NOD-*Prkdc^scid^-IL2rg^Tm1^*/Rj) were purchased from Janvier Labs, France. Mice were group housed under specific pathogen-free (SPF) conditions at the biomedical services unit (BSU), University of Liverpool, Liverpool, UK. Male NSG mice used at TRS laboratories were purchased from The Jackson Laboratory, USA, and group housed within filter-top cages. Mice were age 5-7 weeks and weighed 21-32 g at the start of experiments. Animals had continuous access to fresh sterile food and water throughout experiments. Weight was monitored twice weekly and welfare behaviour monitored daily. Study protocols were approved in the UK by LSTM & University of Liverpool Animal Welfare and Ethics Review Boards and licensed by The UK Home Office Animals in Science Regulation Unit. In the USA, studies were approved by TRS Institutional Animal Care and Use Committee.

### Dirofilaria immitis parasite production

Missouri isolate (MO) *Dirofilaria immitis* microfilariae in dog blood were fed to female *Aedes aegypti* mosquitoes (Liverpool strain) at a density of 5000 mf/ml through an artificial membrane feeder (Hemotek, UK). Blood-fed mosquitoes were reared for 15 days with daily sugar-water feeding to allow development to the L3 stage. At day 15, *Di*L3 were collected from infected mosquitoes by crushing and concentration using a Baermann’s apparatus and Rosewell Park Memorial Institute (RPMI) 1640 with 1% penicillin-streptomycin (both Sigma-Aldrich, UK). For validation studies at TRS Labs, USA, and in-house Georgia III (GAIII) isolate of *D. immitis* was utilised. *Dirofilaria immitis* mf were used to infect female *Aedes aegypti* mosquitoes (Liverpool strain) in dog blood using a glass feeder at a density of 1000 – 2500 mf/ml. At day 14, *Di*L3 were collected from infected mosquitoes using crushing and straining with RPMI 1640 and 1% penicillin-streptomycin.

### Dirofilaria immitis experimental infections

Highly motile infectious stage larvae (*Di*L3) retrieved from mosquitoes were washed in RPMI 1640 with 1% penicillin-streptomycin and 1% amphotericin B (Sigma-Aldrich, UK), and injected subcutaneously into the flank of male NSG/NXG or RAG2ɣc mice at a density of 200 *Di*L3 per mouse. Cohorts of mice also received a single intraperitoneal injection of 2 mg methylprednisolone acetate (MPA; Sigma-Aldrich, UK) immediately prior to infection and after one week post-infection. Mice were humanely culled between 14-28 days post-infection. To retrieve parasites, skins were removed and subcutaneous tissue scarified with a sharp scalpel blade. Muscle tissues were similarly scarified. Visceral organs were dissected and viscera, skin (pellet side-up), muscle tissues and carcass soaked in warm Eagle’s minimum essential media (EMEM; Sigma-Aldrich, UK) with 1% penicillin-streptomycin and 1% amphotericin B for 2 hours to allow active larvae to migrate from tissues. Skin, muscle and carcasses were incubated for a further 24-hour period allowing residual larvae to migrate out of tissues.

### In vitro larval cultures

Madin-Darby Canine Kidney (MDCK) cells and Rhesus Monkey Kidney Epithelial Cells (LLCMK2) cells were passaged in T-75 flasks in EMEM with 10% foetal bovine serum (FBS), 1% penicillin-streptomycin, 1% amphotericin B and 1% non-essential amino acid solution (NEAAS) (Sigma-Aldrich, UK). Cells were seeded onto 12-well plates to reach confluent monolayers 24-48 hours prior to parasite addition. For parasite cultures, washed MO *Di*L3 from mosquitoes were plated onto cell monolayers, or the cell-free media (EMEM) control at a density of 10-20 iL3 per well with 4 ml media. Larvae were monitored over a 35-day time-point for survival and motility and at 14 days post-culture to evaluate development, length and *Wolbachia* titres.

### In vitro and ex vivo drug screening assays

MO *Di*L3 larvae were transferred onto MDCK monolayers and allowed to develop to 14 (early-mid L4) or 28 (mid-late L4) day old larvae. For comparative *ex vivo* assays, L4 stage larvae were recovered from male NSG mice 14-days post infection and washed in sterile EMEM prior to the addition of drugs. All larval stages were plated into 12-well plates at densities of 3-5 larvae per well per drug concentration in 4ml of EMEM with 10% FBS, 1% penicillin-streptomycin, 1% NEAAS and 1% amphotericin B for drug screening assays. Moxidectin (Sigma-Aldrich, UK) was solubilised in phosphate buffered saline (PBS, Fisher Scientific) and 10-fold serial dilutions ranging from 0.0001–100µM were prepared in EMEM with 1% penicillin-streptomycin, 1% NEAAS and 1% amphotericin B. Vehicle controls were included using the equivalent percentage PBS added to the cultures. Assays were incubated for 6 days in which larvae were continuously exposed to drug at 37°C, 5% CO_2_, and scored daily for motility and survival.

### In vivo drug screening validation

Paired groups of 1-5 male NSG mice were subcutaneously inoculated with 200 *Di*L3 into the right flank on day 0. They were then randomised into treatment groups with a single subcutaneous dose of moxidectin prepared at 2.5 mg/kg in saline, or a saline only control, in the nape of the neck on day 1. Mice were monitored daily for weight change and culled at day 14 to evaluate efficacy based on parasite recoveries. Alternatively, immediately following infection, groups of 4-6 mice were randomised into a 7-day oral regimen of doxycycline at 50 mg/kg prepared in ddH_2_O followed by a 7-day washout period, a 2-day oral bi-daily regime of AWZ1066S prepared in standard suspension vehicle (SSV; PEG300/propylene glycol/H_2_O (55/25/20), or matching vehicle controls. Mice were monitored daily (weight and welfare) and culled between day 14-28 post-infection to evaluate parasitology, L4 length, and/or *Wolbachia* depletion using qPCR.

### Wolbachia titre analysis

To assess *Wolbachia* titres across different developmental timepoints (2-, 3- and 4-weeks post infection), to compare *in vitro* reared larvae in parallel to *in vivo* reared controls and to investigate drug activity against *Wolbachia*, individual larvae were taken, and their DNA extracted using previously published methods [15]. *Wolbachia* single copy *Wolbachia surface protein* (*wsp*) gene quantification was undertaken by qPCR using the following primer pair: F-TTGGTATTGGTGTTGGCGCA and R-AGCCAAAATAGCGAGCTCCA, under conditions used to determine *Brugia malayi wsp* copy numbers [15].

### Fluorescence in-situ hybridisation

Fluorescence *in-situ* hybridisation (FISH) was used for detecting *Wolbachia in Di*L3 and L4 larvae using two different DNA probes specific for *Wolbachia* 16S rRNA: W1 -/5ATTO590N/AATCCGGC-GARCCGACCC and W2 -/5ATTO590N/CTTCTGTGAGTACCGTCATTATC, as previously described Walker, Quek (60). L3 and L4 larvae were stored in 50% ethanol at room temperature until further processing. For FISH staining, frozen larvae were fixed using 4% paraformaldehyde (PFA) and incubated with 10ug/ml pepsin for 10 min at 37°C. After thorough wash using PBS, samples were hybridised overnight in hybridisation buffer with probes (or without probes for negative controls). Hybridisation buffer consisted of 50% formamide, 5xSCC, 0.1M dithiothreitol (DTT), 200g/L dextran sulphate, 250mg/L poly(A), 250mg/L salmon sperm DNA, 250mg/L tRNA and 0.5x Denhardt’s solution. Larvae were then washed twice in 1xSSC and 0.1xSSC 10Mm DDT before mounting with VECTASHIELD antifade mounting medium containing DAPI (4’,6-diamindio-2-phenylinole) (Vector laboratories). L4 were visualised using brightfield microscopy for length measurements, calculated using Fiji (ImageJ), USA. FISH-stained larvae were imaged using Zeiss laser scanning confocal microscope and changes in larval morphology was visualized using brightfield and DAPI nuclear staining.

### Statistical analysis

Continuous data were tested for normality using the using D’Agostino & Pearson omnibus Shapiro–Wilk normality tests. Where data were skewed, non-parametric analyses were used to compare statistical differences between groups using Dunn’s post-hoc tests. Where data passed normality tests, Tukey’s post-hoc tests were applied. Categorical data was analysed by Fisher’s Exact Tests. Survival of larvae in culture (frequency motile vs immotile) was evaluated by Log-rank (Mantel-Cox) test. Moxidectin / levamisole IC_50_ values were derived from percentage immotile larvae per drug concentration on day 6 of the assay. Non-linear curves were generated by 3-parameter least squares fit with [IC_50_] calculated. All tests were performed in GraphPad Prism 9.1.2 software. Significance is indicated at or below alpha = 0.05.

## ACKNOWLEDGEMENTS

We gratefully acknowledge the NIH/NIAID Filariasis Research Resource Center (www.filariasiscenter.org) for donation of the *D. immitis* Missouri 2005 isolate.

## AUTHOR CONTRIBUTIONS

Conceptualization: Joseph D. Turner. Data curation: Amy E. Marriott, Jessica L. Dagley. Formal analysis: Amy E. Marriott, Jessica L. Dagley, Shrilaksmi Hegde, Joseph D. Turner. Funding acquisition: Mark J. Taylor, Joseph D. Turner. Investigation: Amy E. Marriott, Jessica L. Dagley, Shrilaksmi Hegde, Andrew Steven, Crystal Fricks, Utami DiCosty, Abdelmoneim Mansour, Caro M. Wilson, Elyssa J. Campbell. Methodology: Amy E. Marriott, Jessica L. Dagley, Shrilakshmi Hegde. Project administration: Joseph D. Turner, Andrew Moorhead, Scott McCall, John W. McCall. Resources: Fabian Gusovsky, Steven Ward, David Hong, Paul O’Neill, Mark J. Taylor, Joseph D. Turner. Writing – original draft: Amy E. Marriott, Jessica L. Dagley, Joseph D. Turner. Writing – review & editing: Amy E. Marriott, Jessica L. Dagley, Crystal Fricks, John W. McCall, Scott McCall, Andrew Moorhead, Joseph D. Turner.

